# Whole mitogenome analysis highlights demographic history and shared connections among distal Indigenous groups of Mexico Complete mitogenome sequencing from 60 Mexican Native American groups

**DOI:** 10.1101/2023.09.03.556146

**Authors:** Marlen Flores-Huacuja, Meradeth Snow, Jazmín Ramos-Madrigal, Cecilia Contreras-Cubas, Francisco Barajas-Olmos, Angélica González-Oliver, Elvia Mendoza-Caamal, Isabel Cicerón-Arellano, Federico Centeno-Cruz, Emilio J. Córdova, Paulina Baca, Silvia Esperanza Flores-Martínez, Rocío Ortiz-López, Austin W. Reynolds, Aleksandar David Kostic, José Rafael Villafan-Bernal, Carlos Galaviz-Hernández, Alejandra Guadalupe García-Zapién, Haydee Miranda-Ortíz, Blanca Patricia Lazalde-Ramos, Francisco Loeza-Becerra, Alessandra Carnevale, Héctor Rangel-Villalobos, Martha Sosa-Macías, Augusto Rojas-Martinez, Angélica Martínez-Hernández, Humberto García-Ortiz, Lorena Orozco

## Abstract

The study of mitochondrial DNA is a valuable tool to delve into the demographic history of human populations. Particularly in the Americas, five widespread Native American specific mitochondrial lineages have been identified. Here we included the complete mitogenome sequencing of 572 Indigenous individuals belonging to 60 populations spanning the Mexican territory. Our results show a great diversity of matrilineages widespread across the country, revealing shared mtDNA haplogroups in populations from distant regions. We identified all the five main Native American haplogroups clades, including 83 different subhaplogroups, from which nine are novel. The most frequent of the novel haplogroups was A2+64. A phylogenetic inference suggests that A2+64 comes from an ancestral maternal lineage that spread into the Caribbean islands. Additionally, a demographic reconstruction from whole mitogenomes showed an exponential increase in female Ne around 10 Ka ago in all the tested regions. All these findings suggest a genetic persistence through Mexico and possibly the Americas, in agreement with the model of the Mesoamerican-related expansion into the Caribbean and South America.

## Introduction

Mitochondrial DNA (mtDNA) is an extranuclear 16.6 kb double-stranded DNA encoding for 37 genes. It is maternally inherited without recombination, possesses a higher mutational rate in comparison with the nuclear genome and it has a high copy number per cell [1]. Due to a process of random genetic drift and selection, several specific mtDNA lineages are widespread and fixed among human populations, making it a valuable tool to elucidate migration routes, human populations distribution patterns and evolutionary reconstructions [1,2].

The study of Native American mitochondrial lineages has contributed to elucidate how the complex demographic history, repeated and rapid populations movements throughout the continent, bottlenecks, and social conflicts have shaped the genetic structure of past and present day Native American populations [3]. To date, six specific Native American mitochondrial haplogroups (A2, B2, C1, D1, D4h3a and X2a) distributed in the American continent have been described. It has been suggested that these matrilineages come from a founding population that split from Northern Eurasian lineages around 25 ka ago and remained stranded in Beringia for ∼6 ka before they moved into the Americas ∼16 ka ago [4–7]. The distribution and frequency of these lineages varies across the continent and within local populations [8].

To better understand the distribution of these mitochondrial haplogroups, most investigations have focused on studying mtDNA hypervariable I & II regions, or RFLP haplogroups [9–18]. These studies have also identified specific regional admixture patterns and high overall haplogroup/haplotype diversity between populations, mainly between those from the North (Aridoamerica), and those from the Center and South (Mesoamerica) of Mexico [15,17,19].

Recently, to deepen our understanding in the peopling of the Americas, whole ancient and modern mitogenomes have been utilized [6]. These studies have demonstrated that the demographic bottleneck that the first settlers experienced, along with their rapid expansion through the continent and limited intracontinental geneflow, have resulted in a marked phylogeographic populations structure, which has persisted through time [3–6]. It has been suggested that the first wave of people entering into the current Mexican territory brought with them the A2 and B2 haplogroups around 15-18 ka ago, followed by a series of expansions that brought additional diversity with the haplogroups C1 and D1 [20]. Previous studies analyzing whole mitogenomes from Mayans, Mazahuas, and Zapotecs, all of them Mexican Native Americans, suggested two main paleo-groupings: i) Centro-Mesoamerican which remained within Mesoamerica, and ii) Pan-American, which moved to South America [19,21].

Although mtDNA studies have addressed the genetic structure and migration of Mexican Native Americans, most of them have often been limited in scope and sampling [3]. The Mexican territory encompasses 68 Native American populations with unique cultural and demographic histories, such as the establishment of sedentary agriculture in Mesoamerica, or the European contact [22,23]. To gain insights into the evolutionary history of the mtDNA lineages in Mexican population, we analyzed the complete mitogenomes of 572 Native American individuals representing 60 populations spanning the country. The significant increase in the number of whole mitogenomes sequenced and number of Native American groups presented here, along with ancient mitogenomes from the Americas make it possible to look in greater detail at the regional variation present in Mexico and too deep in the genetic relationships of these groups, identifying novel mitochondrial subhaplogroups as well as shared matrilineages that span the entire country, helping to delve into the genetic differences between populations that belong to the same linguistic family. These findings are indicative of either shared ancestral maternal lineages or potentially population movements due to different demographic phenomena. The current data also brings insights into the genetic and cultural exchange that have taken place in the Americas through time. This is the first study that incorporates a large proportion of mitogenomes in a great diversity of Native American populations.

## Results

### Variants Call and Haplogroup Identification

To minimize the impact of admixture, especially with Europeans, we only included individuals with a Native American ancestry greater than or equal to 90%, estimated with ADMIXTURE v1.30 [24] (S1 Fig). Mitogenome sequencing was carried out in a total of 572 Mexican Native Americans encompassing 60 populations from 75 indigenous communities belonging to the Metabolic Analysis in an Indigenous Sample (MAIS) cohort (S1 Table). Raw reads were mapped to the Cambridge reference sequence for human mitochondrial DNA [25] and variant discovery was performed using HaplotypeCaller from GATK v3.7, setting the ploidy parameter to 1 [26,27]. Based on this analysis, 952 variants passed the quality filters: 923 SNVs and 29 indels.

### Distribution and frequency of mitochondrial haplogroups

Subsequently, the identification of haplogroups was carried out using Haplogrep2 v2.1.18 [28,29]. Three non-Native American haplogroups were identified in four individuals, which were excluded from the subsequent analyses (S2 Table). In the final data set of 568 mitogenomes, we identified five Native American haplogroups A2, B2, C1, D1 and D4h3a, where the most prevalent was A2 (53.9%), followed by B2 (24.0%), C1 (17.6%), D1 (4.3%) and D4h3a (0.2%). This general pattern follows previous findings of haplogroup frequencies in Mexico [15,16,30].

To get a deeper insight into the haplogroups distribution, we compared their geographic distribution using only those population with a sample size greater than or equal to 5 individuals (532 mitogenomes distributed in 52 indigenous populations). Although the distribution of the four main haplogroups (A2, B2, C1, D1) was highly heterogeneous throughout the Mexican territory, they were present in all regions of the country, with different frequencies among regions and populations (Fig 1). For example, we observed the highest frequency of A2 haplogroup in populations from the Southeast, Center and South of Mexico (Table 1) and notably, Huave (South, n = 10), Lacandon (Southeast, n = 11) and Kaqchikel individuals (Southeast, n = 10) only carried this haplogroup (Fig 1 and S3 Table). Moreover, the B2 and D1 haplogroups were more frequent in the Northwest (40.5% and 13.5%, respectively), while the C1 in the North (36.6 %) (Table 1). We also identified that haplogroup B2 exhibits higher frequency than A2 in eight populations from four linguistic families (Wixarika, Mexicanero, Tepehuano, Mazahua, Mazateco Puebla, Totonaco Puebla, Zapoteco and Mocho). While, the haplogroup C1 exhibits higher frequency than A2 in Seri, Purepecha and Raramuri (Fig 1 and S3 Table).

**Fig 1.**
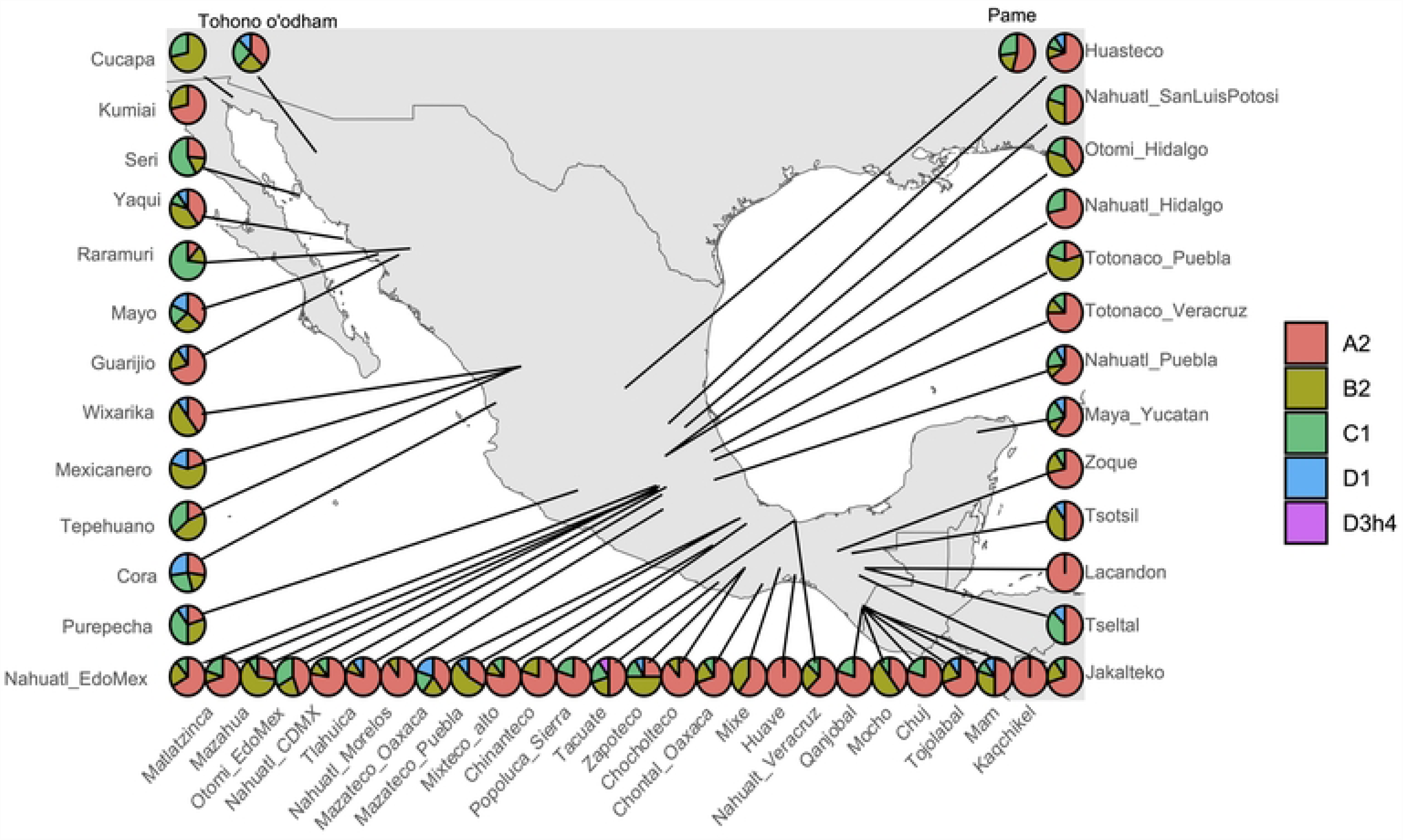
Haplogroup distribution in 52 indigenous populations. The points in the graph show the approximate location and the pie charts show the proportion of the main haplogroups in each studied population.

**Table 1.** Haplogroups count and frequencies by geographic region (n and %).

| Region | n | A2 |  | B2 |  | C1 |  | D1 |  | D4h3 |  |
| --- | --- | --- | --- | --- | --- | --- | --- | --- | --- | --- | --- |
| North | 101 | 32 | 31.7 | 27 | 26.7 | 37 | 36.6 | 5 | 5.0 | 0 | 0.0 |
| Northwest | 37 | 10 | 27.0 | 15 | 40.5 | 7 | 18.9 | 5 | 13.5 | 0 | 0.0 |
| Center | 185 | 112 | 60.5 | 38 | 20.5 | 31 | 16.8 | 4 | 2.2 | 0 | 0.0 |
| South | 92 | 53 | 57.6 | 25 | 27.2 | 9 | 9.8 | 4 | 4.3 | 1 | 1.1 |
| Southeast | 153 | 103 | 67.3 | 29 | 19.0 | 15 | 9.8 | 6 | 3.9 | 0 | 0.0 |

On the other hand, 82 mitochondrial subhaplogroups were identified, 34 related to A2, 29 to B2, 12 to C1, 7 to D1, all of them with a variable frequency ranging from 0.2-5.6 % for all branches (Table 2). Notably, we found nine subhaplogroups non-previously reported in the present-day Native American populations, five related to the A2 haplogroup, three derived from B2 haplogroup, and one from the C1 haplogroup (Table 2, bold text). Moreover, 50 additional subhaplogroups, previously reported in other Native American populations across the continent, were also identified for the first time in the examined Mexican Native American individuals. All of them had a low frequency, ranging from 0.2% to 5.6%, among them A2+64 showed the highest frequency (5.6%), mainly in those populations related to Mixe-Zoque (19.5%, n = 41) and Mayan linguistic families (18.9%, n=132), both from the Southeast region (S4 Table). To investigate the A2+64 divergence time, we analyzed 50 Mexican Native American individuals and seven ancient samples from Puerto Rico previously reported as carrying the A2+64 haplogroup [31]. To calibrate the tree, we included eight ancient samples and five present-day individuals harboring other haplogroups. The Bayesian species calibrated tree inferred under a strict molecular clock showed that the ancient and modern A2+64 lineages fall in the same clade, suggesting the same origin for both samples (Fig 2, S2 Fig). Moreover, tip dating estimation showed that the origin of A2+64 mitochondrial lineage is between 20 and 10 ka (mean = 14,100 years). These times broadly overlap with the estimated entrance of modern humans into the Americas, suggesting that this lineage was originated with the entrance of the first settlers to the continent.

**Fig 2.**
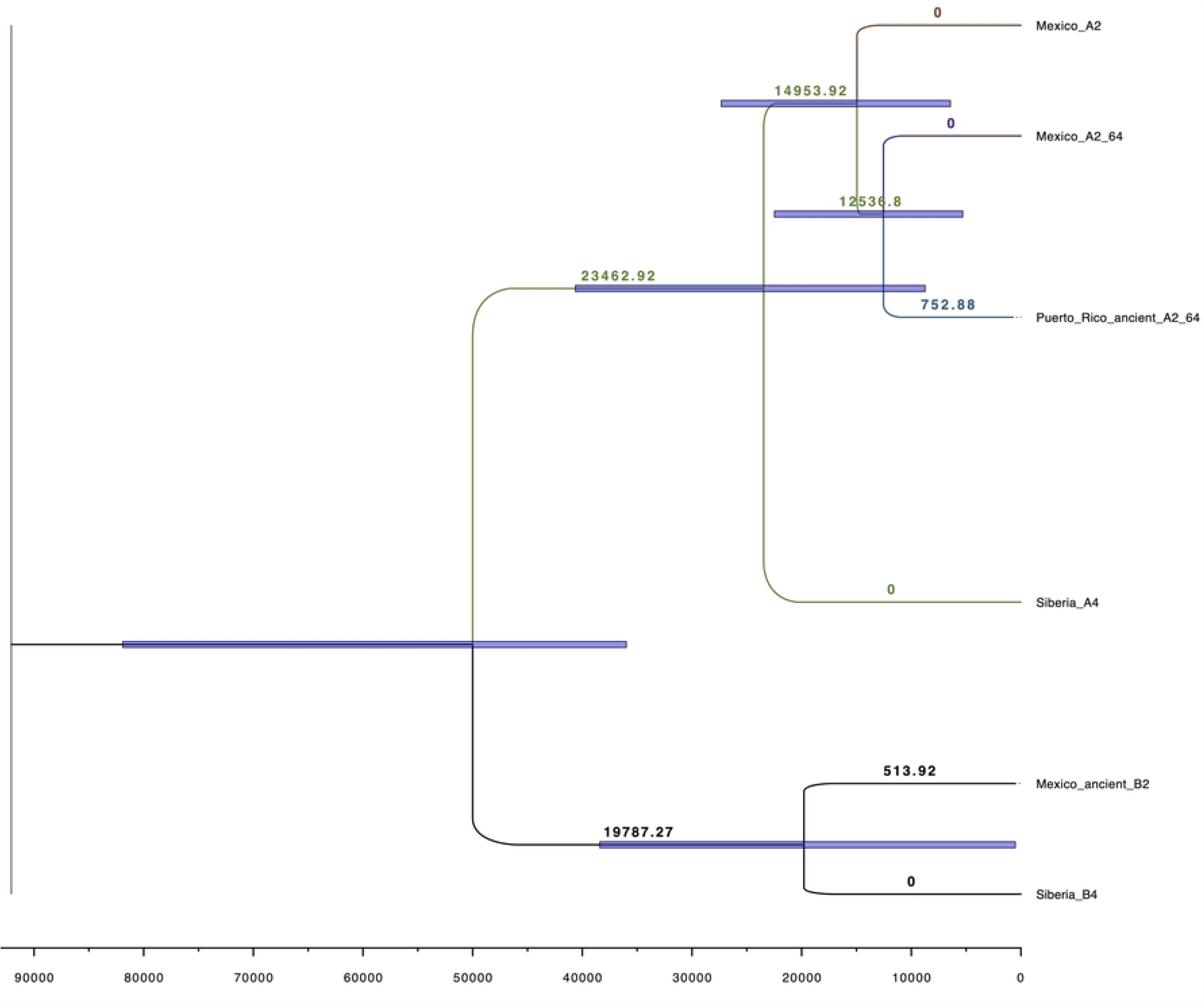
A2+64 haplogroup tip dating tree. A phylogenetic tree was constructed with the whole mitogenome sequences of modern A2+64 carriers using a Bayesian approach under a strict clock model. We included ancient genomes as calibrating information for the tree. Purple bar plots illustrate the 95% HPD values of estimated mean ages for the diversification of A2+64 mitogenomes.

**Table 2.** Haplogroup diversity in the data set (Haplogroup, global frequency).

| A2 |  | B2 |  | C1 |  | D1 |  | D4h3a |  |
| --- | --- | --- | --- | --- | --- | --- | --- | --- | --- |
| A2 | 4.8 | B2 | 4.8 | C1 | 5.3 | D1 | 1.8 | D4h3a | 0.2 |
| A2+(64) | 5.6 | B2+16278 | 1.4 | C1b10 | 0.2 | D1d2 | 0.2 |  |  |
| A2+(64)+@153 | 1.6 | B2+16279 | 0.2 | C1b5a | 0.9 | D1f3 | 0.5 |  |  |
| A2+(64)+@16111 | 1.4 | B2a | 0.5 | C1c | 1.2 | D1h1 | 0.4 |  |  |
| A2+(64)+@16112 | 0.2 | B2a1 | 0.4 | C1c1a | 2.5 | D1i2 | 0.5 |  |  |
| A2ae | 0.4 | B2a1a1 | 0.2 | C1c1b | 0.7 | D1k | 0.2 |  |  |
| A2af1b | 0.7 | B2a4 | 0.4 | C1c2 | 0.4 | D1m | 0.7 |  |  |
| A2ak | 0.2 | B2a4a | 0.2 | C1c4 | 1.1 |  |  |  |  |
| A2ao | 0.4 | B2a4a1 | 1.1 | C1c5 | 2.1 |  |  |  |  |
| A2c | 3.2 | B2a5 | 0.4 | C1c6 | 0.2 |  |  |  |  |
| A2d | 3.0 | B2b | 0.9 | C1d+194 | 0.2 |  |  |  |  |
| A2f2 | 0.4 | B2b+152 | 0.2 | C1d1 | 2.8 |  |  |  |  |
| A2g | 3.0 | B2b2 | 0.4 |  |  |  |  |  |  |
| A2h1 | 1.1 | B2c | 0.7 |  |  |  |  |  |  |
| A2j | 3.0 | B2c1 | 1.6 |  |  |  |  |  |  |
| A2j1 | 0.2 | B2c2 | 1.2 |  |  |  |  |  |  |
| A2l | 0.2 | B2c2a | 0.9 |  |  |  |  |  |  |
| A2m | 4.0 | B2c2b | 0.2 |  |  |  |  |  |  |
| A2o | 0.9 | B2f | 0.7 |  |  |  |  |  |  |
| A2p | 1.6 | B2g1 | 0.5 |  |  |  |  |  |  |
| A2q | 1.8 | B2g2 | 0.2 |  |  |  |  |  |  |
| A2q1 | 0.5 | B2k | 0.4 |  |  |  |  |  |  |
| A2r | 2.0 | B2l | 0.7 |  |  |  |  |  |  |
| A2r1 | 1.1 | B2o | 0.9 |  |  |  |  |  |  |
| A2s | 0.5 | B2s | 0.2 |  |  |  |  |  |  |
| A2t | 1.6 | B2t | 2.5 |  |  |  |  |  |  |
| A2u | 2.3 | B2w | 0.5 |  |  |  |  |  |  |
| A2u1 | 0.5 | B2x | 1.2 |  |  |  |  |  |  |
| A2u2 | 0.2 | B2y1 | 0.5 |  |  |  |  |  |  |
| A2v1 | 0.4 |  |  |  |  |  |  |  |  |
| A2v1+152 | 2.1 |  |  |  |  |  |  |  |  |
| A2v1a | 0.5 |  |  |  |  |  |  |  |  |
| A2w | 4.1 |  |  |  |  |  |  |  |  |
| A2x | 0.4 |  |  |  |  |  |  |  |  |

### Inferring Evolutionary Relationships

The evolutionary relationship was explored in the 52 populations data set. A Nei’s genetic distances matrix was constructed based on complete mitogenome sequences (S5 Table) and a PCoA projection was used to investigate if the populations grouped according to the North, Northwest, Center, South, and Southeast regions from Mexico as previously described [22,32] or linguistic filiation according to INALI classification [33]. These results showed that the populations do not group by region or linguistic affiliation. In contrast, some northern populations are more similar to those from the Southeast. For example, the Wixarika (Northwest) population is close to the Mocho (Southeast) population, and Guarijio (North) is closest to Tojolabal (Southeast). Otherwise, at regional level, a specific linguistic group as Otomi-Pame from the Oto-mangue family, shows a clear subdivision between the closely related Matlatzinca and Mazahua and the Otomi and Pame. Altogether, suggest that they share a common ancestor or high levels of gene flow have shaped this genetic structure (Fig 3 and S6 Table).

**Fig 3.**
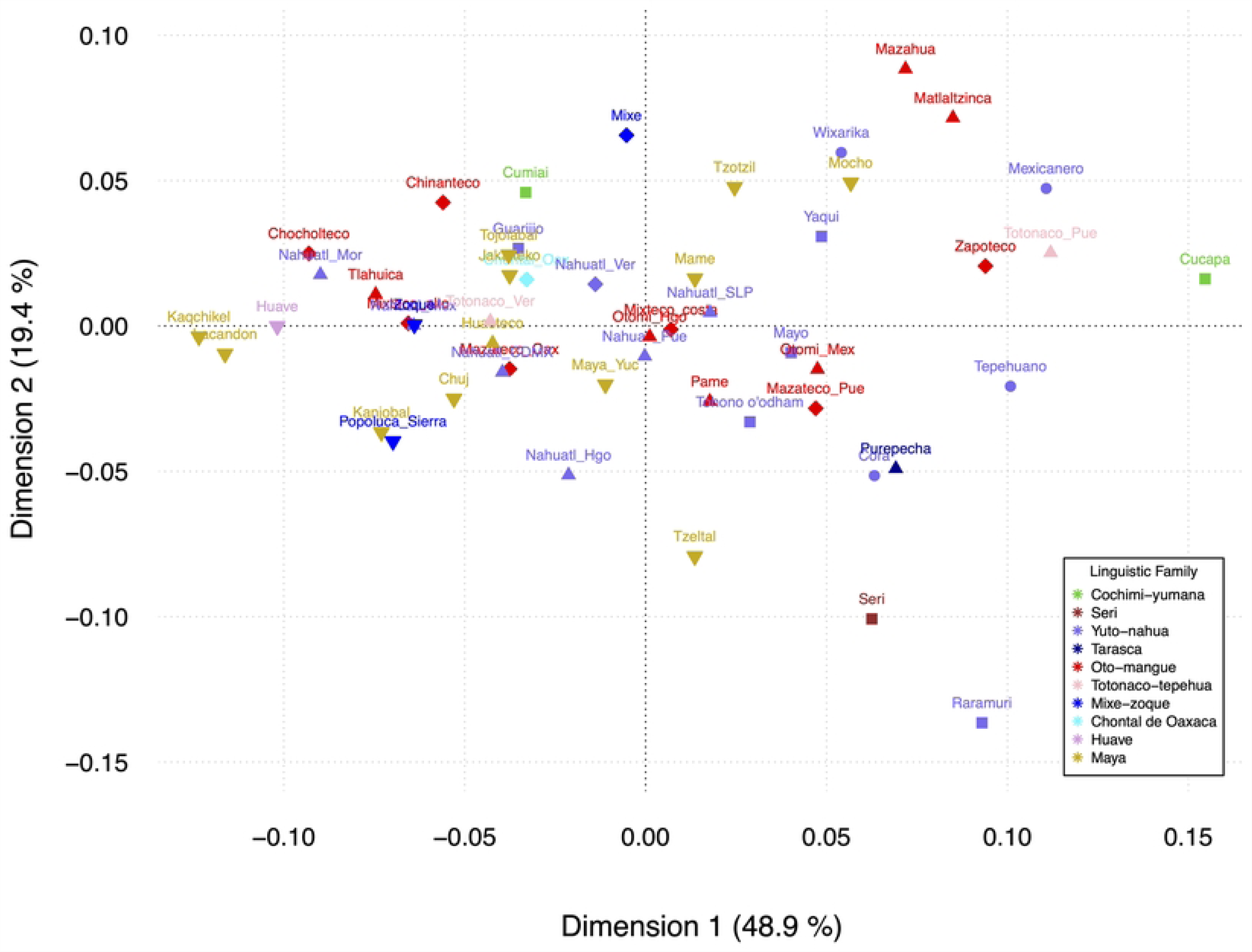
Principal Coordinate Analysis. Graph depicting the matrix of Nei’s genetic distances based on the variable sites of the mitochondrial genome. 52 indigenous populations from Mexico are shown divided by region. Colors indicate the linguistic family and shapes the geographic region in which the populations were classified: North (squares), Northwest (circles), Center (Up-pointing Triangles) South (diamonds) and Southeast (Down-pointing Triangles).

Furthermore, we constructed Median Joining Haplotype Networks for each haplogroup included in the 568 mitogenome data set using PopArt v1.7.2 [34]. These networks showed that samples clustered according to their haplogroup regardless of the geographic region. In general, this pattern is consistent with a prior lineage expansion followed by a local differentiation of the derived haplotypes and pursued by gene flow between different regions (Fig 4). Moreover, a Mantel test showed no correlation between matrilineal population relationships and geographic distance (r^2^ = 0.0167 and p = 0.081; S3 Fig, S5 and S7 Tables).

**Fig 4.**
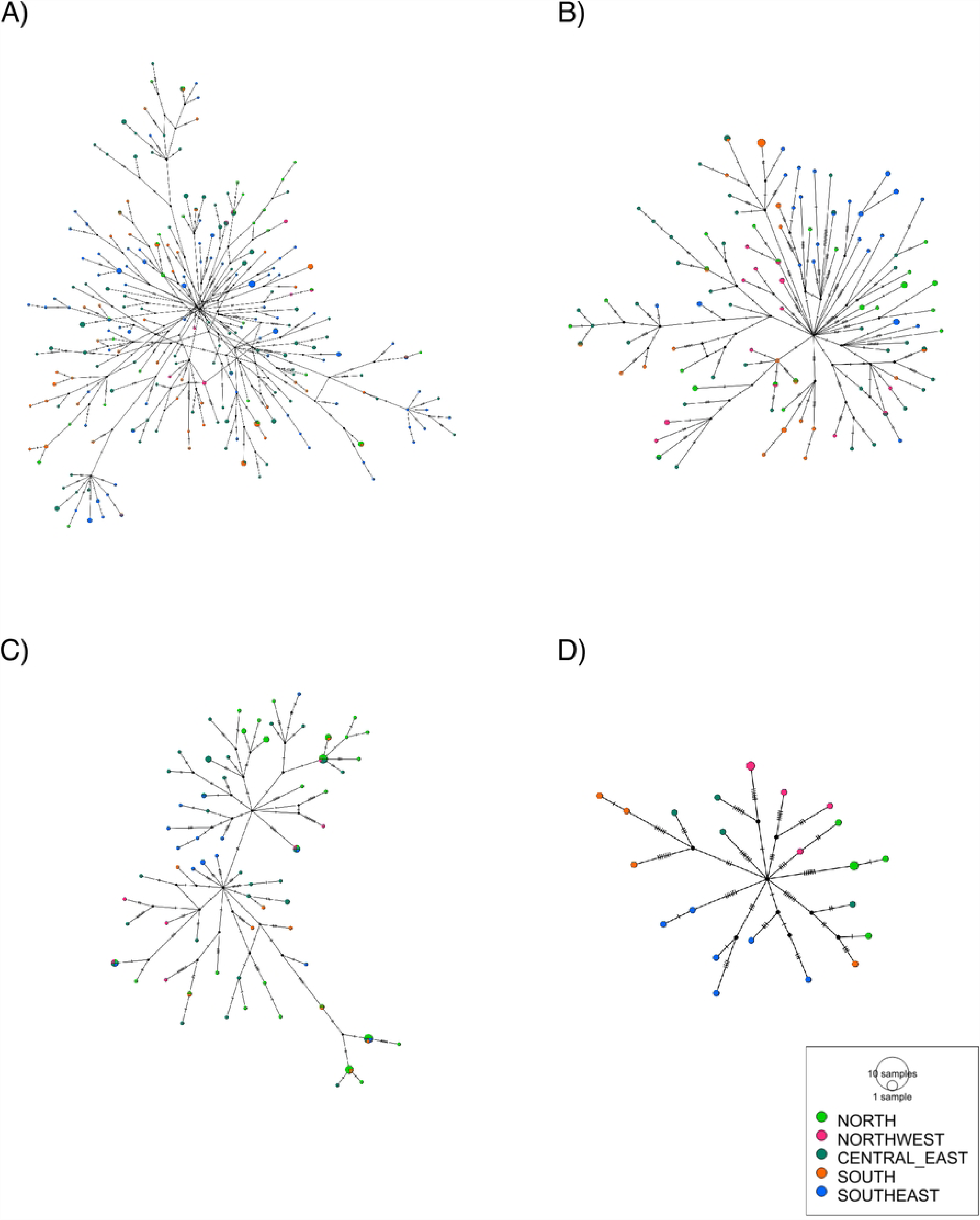
Median Joining Haplotype Networks of whole sequenced mtDNA. (A) A2 haplogroup (B) B2 Haplogroup (C) C1 Haplogroup (D) D1 Haplogroup.

### Estimating Past Population Dynamics

We estimated the past population dynamics and the effective population size (Ne) through time in the set of 568 individuals, using Bayesian Skyline plots (BSP). The runs were broken down by the five main geographical patterns from Mexico. These demographic reconstructions showed a bottleneck resulting in a decline in female Ne in the last 20 generations in all regions, in accordance with previous observations [22,35,36]. Interestingly, the BSP showed an increase of female Ne around 10 ka that was constant until the bottleneck in all the studied regions, suggesting a population expansion of all clades in the early settlement of the present-day Mexican territory (Fig 5).

**Fig 5.**
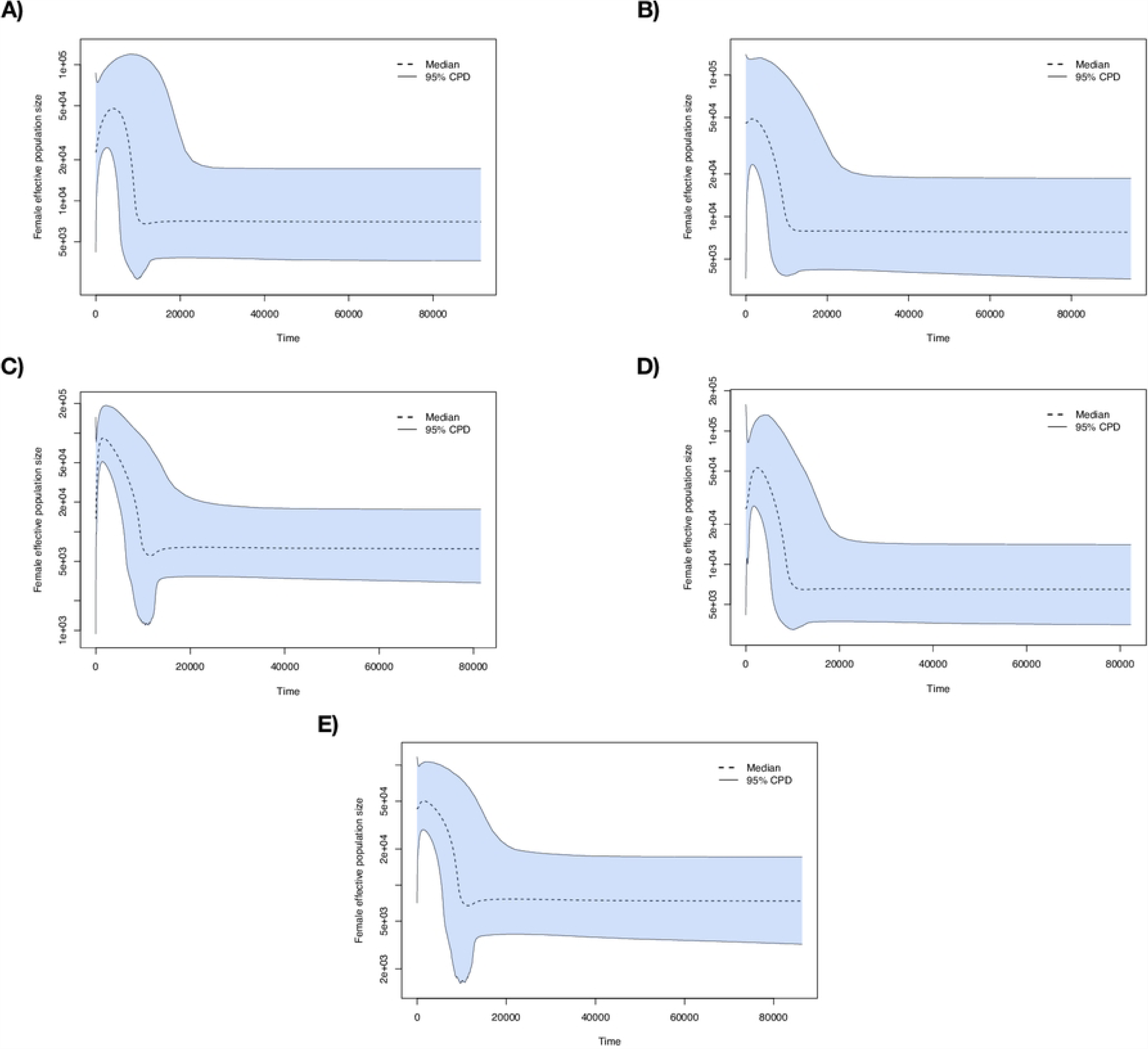
Extended Bayesian skyline plot of female effective population size with calibrating information from ancient sequences. Populations were grouped according to the geographic region. (A) North; (B) Northwest; (C) Center; (D) South; (E) Southeast

## Discussion

Since pre-Hispanic times, different migrations, wars, conquests, territorial removals, and displacements occurred between the numerous Mexican populations. Each of these events translates into bottlenecks, genetic drift, and gene flow that have affected the distribution of the maternal haplogroups of current populations. Several studies have shown that many Mesoamerican populations exhibit the haplogroup pattern ABCD, according to their relative frequency. The extensive geographic sampling included here shows that Native American populations matrilineages have persisted over time with a high haplogroup diversity that vary across Mexico and the Americas. Herein, we identified 5 haplogroups (A2, B2, C1, D1 and D4h3a) following the expected frequency pattern. However, several populations exhibited haplogroups with higher frequency than A2 (Fig 1) The geographic location of individuals showing higher frequencies of B2 or C1 than A2, suggests that A2, B2 and C1 were dispersed mainly by a Pacific coast route and were spread latitudinally 800 km along the Neovolcanic axis region, then southward by the Pacific coast rout and to Yucatan Peninsula via Gulf coast route. Two important rivers, Lerma and Balsas, originate in the Neovolcanic axis which acts as natural barriers to inclement weather from the Pacific, could have also contributed to spreading of the haplogroups (S3 Table).

In addition, 82 different subhaplogroups were also identified, all of them related to the main four haplogroups (A2, B2, C1, D1). From these, 24 subhaplogroups were previously described in the Mexican population, here we report 58 found for the first time in present day Mexican Natives Americans (Table 2). The most frequent among them was the A2+64, which was found with the highest frequency in the populations from the Southeast region belonging to the Mayan (18.9%) and Mixe-Zoque (19.7%) linguistic families. This subhaplogroup was also present in other linguistic families, like the Oto-Mangue (4.1%) from the central region of the country (S4 Table). The tip dating analysis showed that the frequency of this maternal lineage increased in some southern Mexican groups between 28 and 16 ka (Fig 2). This haplogroup was previously identified in Pre-Columbian individuals (7%) from Cholula archaeological site [37] and in 7 of 45 (15.6%) ancient settlers of Puerto Rico before the European conquest [31]. The Bayesian inference showed a common origin from those individuals and our modern samples carrying the A2+64 haplogroup (Fig 2), suggesting that it has an origin in an ancestral lineage related to the early settlement of the Americas that was subsequently dispersed into the Caribbean islands through a coastal and/or marine route. Perhaps, the presence of this subhaplogroup in the Center of Mexico is also due to the gene flow between populations from both regions, as previously documented by anthropological and nuclear DNA studies [22,38–42].

Regarding D4h3a, which is a one of the founding pan-American subhaplogroups, it has been present in the ∼12.8 ka old Anzick-1 child from the Clovis burial and the ∼10 ka old Sumidouro5 from Lagoa Santa, Brazil [43,44]. This haplogroup has been previously found in low frequency in some Native American groups from the Center and South of Mexico and other regions of America, mainly distributed over the Pacific Coast [18,21,45]. Actually, we identified this haplogroup only in a Tacuate female from the Pacific Coast in the Southern region.

In addition, our results show the presence of all four main mitochondrial haplogroup clades without a clear geographic or linguistic structure in all the tested Mexican indigenous groups. Gene flow among populations was evidenced by PCoA and Median Joining Haplotype Networks, in which populations from different regions are closer to each other and share the same matrilineages (Figs 3 and 4). This pattern can be attributed to the constant movements among populations from different regions [22,38]. For example, there is evidence that in prehispanic times, trade contributed to shape the relations between populations, since there were several sites of commercial exchange across the country [38,39,46]. Additionally, the spread of agriculture could have altered the dynamics and the genetic landscape of populations by the dispersal of different haplotypes, mainly B2a5, which emerged approximately 5 ka ago as a lineage derived from B2a [20]. However, at regional level interesting genetic relationships can be found as the subdivision of Oto-mangue speaking populations in Central Mexico in two clearly different groups, one including the Matlatzinca and Mazahua, and the other the Otomi and Pame (Fig 3). The Otomi-Pame linguistic group include only five populations, four were here analyzed. Their linguistic affinity supports a distant common ancestry but separated early in two groups and probably admixed with other populations before migrating to settle in Central Mexico [18]. This genetic difference subsisted despite their geographical proximity due to their marriage customs related with the patrilocal land inheritance and a late urbanization in the region. Another possibility is that the lack of haplogroup structure is also due to shared ancestral lineages, which first entered and spread throughout Mexico before the establishment and differentiation of these populations. In line with this hypothesis, our Bayesian demographic reconstruction shows an exponential increase in female Ne around 10 Ka ago in all regions tested at the same time (Fig 5). This is in accordance with recent studies that show that the early peopling of the Americas was characterized by a rapid dispersal and early diversification of the first settlers through the continent, which in combination with the effect of distance, geographic, and social barriers led to complex population histories [44,47]. Altogether, these data suggest that the indigenous populations were rapidly dispersed through the Mexican territory allowing the distribution of all haplogroups thorough Mexico at different proportions, before linguistic variation was firmly established, and that in conjunction with more recent expansions and contractions of large-scale societies [15], shaped the mitochondrial haplogroups landscape of Mexican Indigenous populations. Otherwise, the presence of ancient Native American haplogroups in the Mexican Indigenous individuals tested here, can be an indicative of genetic persistence of maternal lineages through the continent, in agreement with the model of the Mesoamerican-related expansion into the Caribbean and South America [3,6,7,18,21]. In brief, the mitogenomes reported here fill gaps about the mitochondrial haplogroups distribution, and our demographic inferences showed that the American Indigenous population’s history was marked by admixture between the populations which allowed the spread of the main haplogroups in the region, potentially predating linguistic diversity.

## Materials and Methods

### Sample description

The samples included in this study belong to the Metabolic Analysis in an Indigenous Sample (MAIS) cohort that was described previously [22,32,48,49]. Briefly, the MAIS cohort was collected between 2011-2015, all individuals were self-recognized as Indigenous members of a specific population, and had parents and grandparents born in the same community. This study was designed in accordance with the Declaration of Helsinki and was approved by the local ethics, research and biosafety human committees of the Instituto Nacional de Medicina Genómica (INMEGEN) in Mexico City (protocol numbers: 31/2011/I and 17/2013/I). All participants provided informed written consent. For some participants, informed consent was translated into their native language, and some individuals signed with their fingerprint.

Whole-blood samples were collected by venipuncture in tubes containing EDTA and genomic DNA was extracted using the QIAmp DNA Blood Maxi kit (Qiagen, Valencia CA, USA) according to the manufacturer’s protocol. The DNA concentration was determined by spectrophotometry (Nanodrop 1000 Spectophotometer).

From the MAIS cohort, we included only those Mexican Indigenous individuals with at least 90% of Native-American ancestry determined by ADMIXTURE v1.30 [25], comprising a total of 572 Native Mexicans belonging to 60 populations. To assess this, the samples were previously genotyped with Affymetrix human array 6.0 or Illumina Infinium array. Next, this set was merged with a reference population panel composed of 50 Native Americans, previously identified without evidence of recent admixture with continental populations [22] and with 50 Europeans, and 50 Africans derived from the 1000 Genomes Project Phase 3 [50]. We then ran ADMIXTURE analyses assuming K=3 clusters, including a block relaxation algorithm as the optimization method, and 100 replicates.

### Mitogenome sequencing/Massively Parallel Sequencing on the Ion Torrent PGM

Mitochondrial DNA was amplified by two long-range PCR reactions using two pairs of previously reported primers [51], producing overlapping 8.2 and 8.6 kb amplicons, followed by subsequent fragmentation by sonication to appropriately sized fragments using ShearTM Plus Reagents Kit. The fragments were then ligated to ion barcoded adaptors, and the adaptor-ligated library was then size-selected for 400-500bp length. Finally, the sequencing reaction was carried out in the Ion PGMTM Systems.

The mtDNA data was de-multiplexed and each sample’s raw reads were mapped to the revised Cambridge Reference Sequence (rCRS) [25] with the BWA algorithm [52] and processed with the Genome Analysis Toolkit (GATK) to recalibrate base quality-scores and perform local realignment around known insertion and deletion (indels).

### Variant Calling of mtDNA-Sequence Data/mtDNA Variant Analysis

Target coverage for each sample was computed with the GATK pipeline [26]. Single nucleotide variants (SNVs) and indels were called with the Unified Genotype module of the GATK and filtered to remove SNVs with annotations indicative of technical artifacts (such as strand-bias, low variant call quality, or homopolymer runs). Variant discovery was performed using HaplotypeCaller from GATK v3.7, setting the ploidy parameter to 1 and next we applied hard filters to SNV and indel calls using the GATK’s VariantFiltration [26,27]. Variants were annotated with The Ensembl Variant Effect Predictor (VEP) [53].

### Haplogroups Identification

Aligned sequences were visualized and checked with MEGA7 v7.0.26 software [54] prior haplogroup determination. The assignment of haplogroups for each sample was assessed with the HaploGrep2 v2.1.0 software, which provides an automated way to determine the haplogroup of the mtDNA profiles, based on Phylotree [28]. The classification of haplogroups was based on pre-calculated phylogenetic weights that correspond to the occurrence per position in Phylotree and show the stability of mutation of a variant [28]. Samples with non-Native American haplogroups, were excluded for subsequent analyses (n=4).

### Tip Dating

To date the A2+64 haplogroup we first transformed the genotypes from 50 individuals from MAIS carrying the A2+64 haplogroup into a fasta file using the *FastaAlternateReferenceMaker* command from GATK v3.7 [26,27]. All fasta sequences were merged into a single file and were aligned with MAFFT v7 [55]. Next, each sequence was checked with MEGA7 v7.0.26. These sequences were loaded in PartitionFinder2 v2.1.1 [56,57] to choose the nucleotide substitution model. To calibrate the tree, we included four ancient samples from Mexico carrying the B2 haplogroup, two contemporary sequences from A4, two from Siberia carrying the B4 haplogroup [6] and seven ancient sequences with the A2+64 haplogroup from Puerto Rico [31]. The Bayesian species calibrated tree was constructed in BEAST2 v2.6.6 [58] with the following parameters: a constant population model, a TN93 as nucleotide substitution model, a strict molecular clock, six partitions (position by codon: 1spot, 2stpos, 3stpos; position by gene: tRNA, rRNA, HVR1-HVR2 and ND6) [6], a mutation rate of 8×10^-8^, four chains of the MCMC algorithm for 500 million iterations each, sampling every 500 steps with a burn-in of 25%. Convergence was assessed based on the effective sample sizes estimates (>200) and loglikelihood distribution through the run.

### Distribution of mitochondrial haplogroups

In order to compare the haplogroup distribution in Mexican Native American, we included only those populations with a sample size ≥ 5 giving a n = 532 samples distributed in 52 indigenous populations. Next, a genetic distances matrix based on Nei’s index were constructed to establish approximate relationships among populations through a Principal Coordinate analysis (PCoA) based on a dissimilarity matrix using the haplogroup designations between populations. To test if mitogenome genetic and geographic distance are related a Mantel test. All tests were carried out with the GenAlex v6.5 software [59].

### Median Joining Haplotype Networks

To infer evolutionary relationships, we carried out Median Joining Haplotype Networks with Population Analysis with Reticulate Trees using PopArt software with standard specifications [34] in the set of 568 mitochondrial genomes. For regional comparisons, we included the regional Mexican country division North, Northwest, Center, South and Southeast proposed in Contreras-Cubas et al [32].

### Demographic inference through Bayesian Skyline Plots

For the five regions, estimation of past population dynamics was carried out by Skyline Plots with BEAST2 (Bayesian Evolutionary Analysis by Sampling trees) v2.4.5 [60] in the set of 568 mitochondrial genomes. BEAST2 is based on Monte Carlo Chains Markov (MCMC) method [61,62]. A Coalescent Extended Bayesian Skyline was run using six partitions (position by codon: 1spot, 2stpos, 3stpos; position by gene: tRNA, rRNA, HVR1-HVR2 and ND6), the T93 model with a strict molecular clock and 250 millions of iterations [59], following Heled and Drummond 2008 [62] and Bouckaert et al 2014 [58]. Also, we used the sequence of Anzick-1 and five ancient sequences from Mexico, 20 modern sequences from Siberia [6], and eight Mexican sequences with African haplogroups with the purpose of calibrating the tree. Trees were plotted with R software [63].

## Acknowledgements

Marlen Flores-Huacuja is a doctoral student from the Programa de Doctorado en Ciencias Biomédicas, Universidad Nacional Autónoma de México (UNAM) and has received the CONAHCyT fellowship 596612. This study was supported by the Consejo Nacional de Ciencia y Tecnología, Mexico [233970]. The funder had no role in the study design, data collection and analysis, decision to publish, or preparation of the manuscript.

## Supporting information

**S1 Fig.**
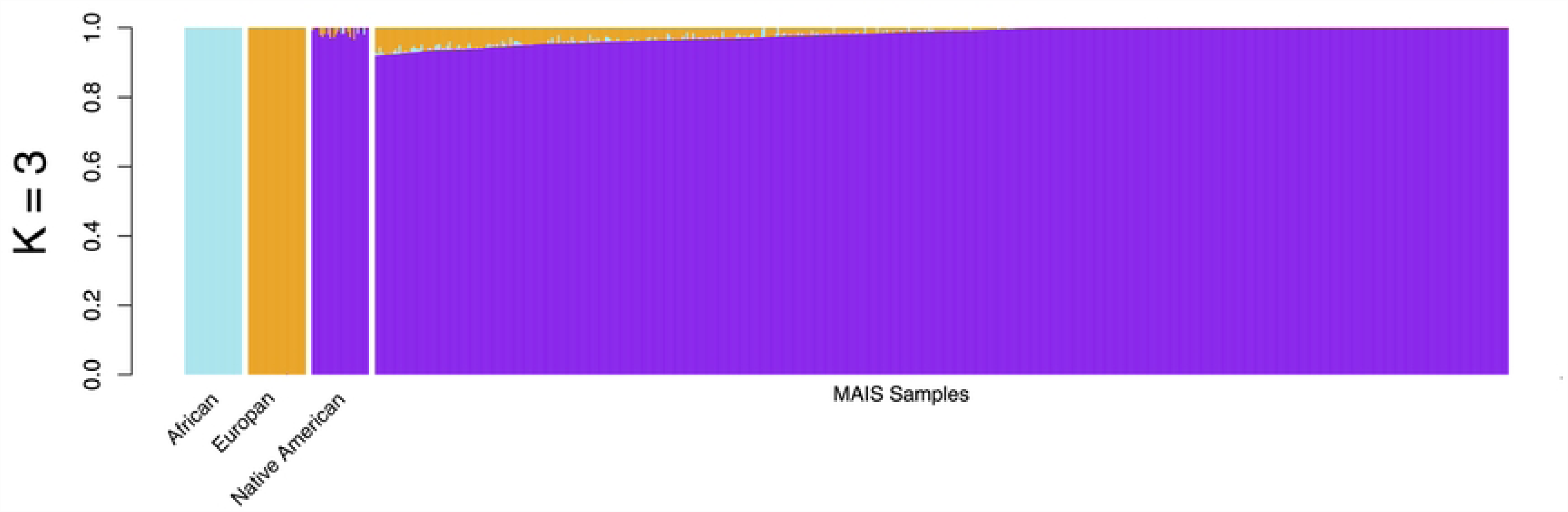
Admixture inference assuming K = 3 clusters. The inference of ancestry proportions in the MAIS dataset was done using a reference panel of African, European and Native-American populations. Purple colored bars represent the proportion of inferred Native American ancestry in each sample.

**S2 Fig.**
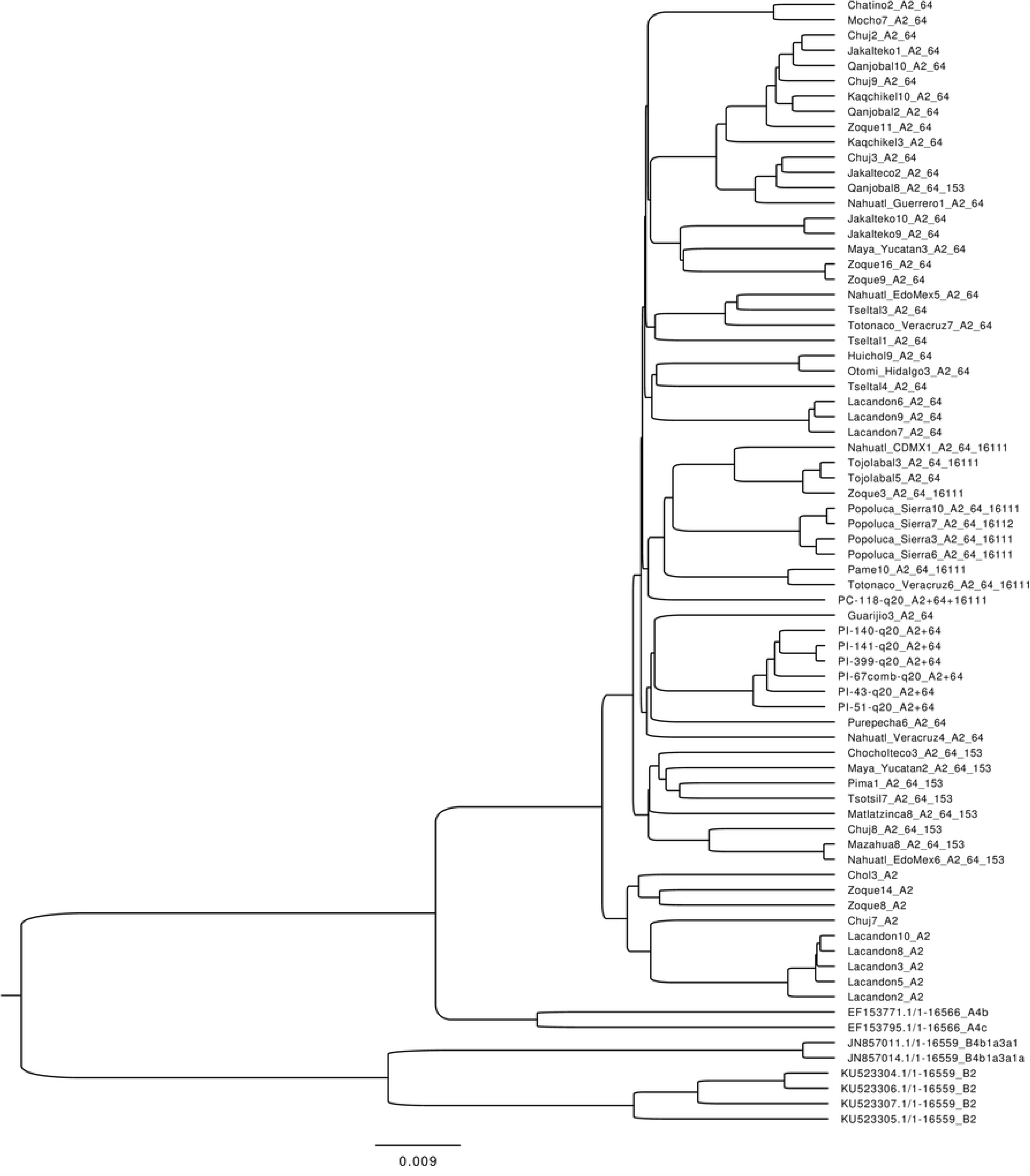
Bayesian phylogenetic analysis of A2+64. Calibrated tree from the whole mitogenome sequences of modern and ancient A2+64 haplogroups mitogenomes

**S3 Fig.**
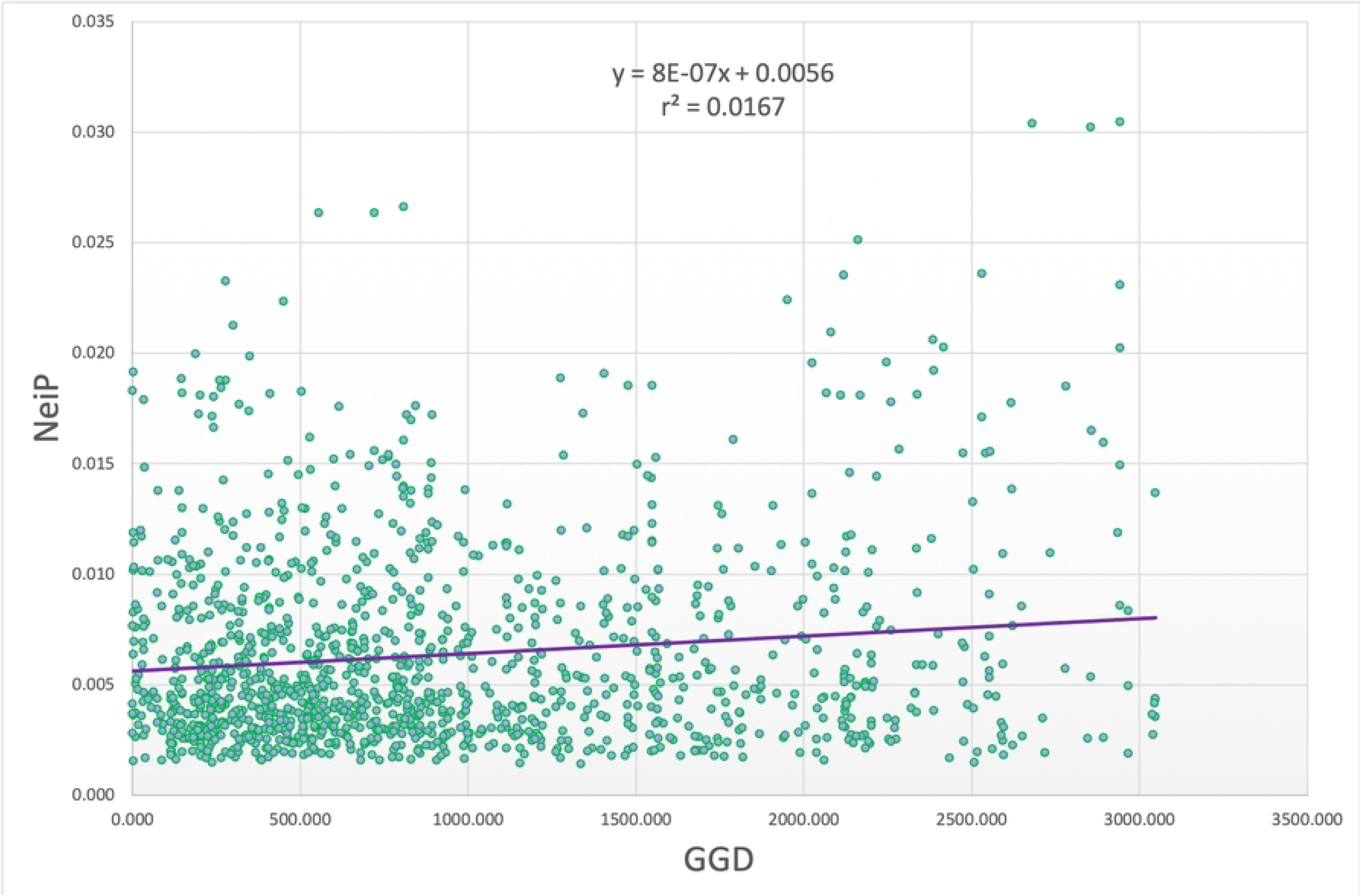
Mantel test. Mantel test with 999 permutations based on Nei’s genetic distances using the variable sites of the mitochondrial genome and geographic distance (GGD).

**S1 Table. Populations size included in the present study.**

**S2 Table. Non-Native American Haplogroups identified in the studied populations.**

**S3 Table. Haplogroup count and frequency by indigenous population (n and %).**

**S4 Table. A2+64 Haplogroup frequency by linguistic Family and ethnic group.**

**S5 Table. Pairwise Nei’s genetic Distance Matrix.**

**S6 Table. Eigen Vectors inferred from Nei’s genetic distance Matrix.**

**S7 Table. Pairwise geographic distance Matrix.**

